# Cell adhesion molecule interaction with Piezo1 channels is a mechanism for sub cellular regulation of mechanical sensitivity

**DOI:** 10.1101/602532

**Authors:** E Chuntharpursat-Bon, OV Povstyan, MJ Ludlow, HJ Gaunt, PD Baxter, DJ Beech

**Affiliations:** School of Medicine, University of Leeds, Leeds, LS2 9JT, UK

## Abstract

The discovery of Piezo1 channels as force sensors with roles in the endothelial response to fluid flow has appeared contradictory to earlier work suggesting mediation by a triad of cell adhesion molecules (CD31 and VE-cadherin) and vascular endothelial growth factor receptor 2 (VEGFR2). Here we propose an explanation. Stimulated emission depletion microscopy revealed physical proximity between Piezo1 and CD31 primarily at cell junctions. Förster resonance energy transfer measured by fluorescence lifetime imaging showed that the proteins approached to less than 10 nm. Substitution of a tyrosine residue at the distal C-terminus of CD31 prevented the interaction, suggesting intracellular association. The extracellular N-terminus also interacted, but exclusively at cell junctions. There was proximity between Piezo1 and VE-cadherin but not VEGFR2. Interaction with VE-cadherin was flow-dependent, consistent with additional recruitment after flow-sensing. CD31 suppressed mechanical sensitivity of Piezo1 channels and interacted with N-terminal Piezo1 propeller arms implicated in force sensing. The data suggest partnership between Piezo1 and adhesion molecules for sub cellular tuning of force response.

## Introduction

Endothelial force sensing is critical in development, health and disease^1–5^. The mechanisms have been studied intensely yet remain unclear, with several apparently contradictory proposals^6–10^. An important hypothesis has involved two adhesion molecules^11^; these proteins are CD31 (a.k.a. platelet endothelial cell adhesion molecule 1, PECAM-1) and vascular endothelial cadherin (VE-cadherin, a.k.a. cadherin 5), which are single-pass membrane proteins that mediate cell contact and junctional integrity^12–14^. They are well positioned for participation in force sensing between cells. CD31 is a 130 kDa protein belonging to type-I membrane glycoprotein and immunoglobulin super-families. It has an extracellular region containing immunoglobulin-like domains, a transmembrane region of one α helix and a cytoplasmic region containing tyrosine regulatory motifs. It is often used as an endothelial cell marker but is expressed and functional also in leukocytes and platelets^12^. VE-cadherin belongs to a large family of transmembrane Ca^2+^-dependent adhesion molecules^14,15^. It also has an extracellular domain that mediates homotypic interactions and an intracellular regulatory domain, which in this case binds β- or γ-catenin to promote cytoskeletal interaction^14^. An influential study linked CD31 and VE-cadherin to sensing of mechanical force caused by fluid flow and suggested that vascular endothelial growth factor receptor 2 (VEGFR2) was recruited as a third partner, leading to the prominent hypothesis that a CD31/VE-Cadherin/VEGFR2 triad is a key flow sensor^8,11,16^. Subsequently another endothelial flow sensing hypothesis emerged because of the discovery of a different protein called Piezo1^17–22^. Piezo1 is a 38-pass membrane protein of 286 kDa that assembles as a trimer to form Ca^2+^-permeable non-selective cationic channels^23–26^. This channel is exquisitely sensitive to membrane tension, activating rapidly as membrane tension rises^26–28^. Importantly it is also activated by fluid flow^17–22^, strongly but not uniquely expressed in endothelial cells^17,28^ and required for endothelial responses to force such as cell alignment in the direction of flow and activation of endothelial nitric oxide synthase^17,20^. It is also implicated in cell junction integrity^29^. Here we investigated the relationship of the CD31/VE-cadherin/VEGFR2 triad to Piezo1.

## Results

To investigate relationships of Piezo1 to other proteins we expressed tagged constructs in the COS-7 immortalised cell line because the flat nature of these cells in culture facilitates high-resolution imaging and they do not natively express, or express very low levels, of the proteins of interest, providing a null or almost null background. First we co-expressed CD31 with Piezo1 engineered to contain a haemagglutinin (HA) tag in its C-terminal extracellular domain (CED). The CD31 was labelled with antibody targeted to a native extracellular N-terminal epitope and Piezo1 with antibody targeted to HA (Figure 1). Cells were unpermeabilised so that labelling was restricted to surface-localised proteins. Confocal imaging revealed enrichment at cell-cell junctions (Figure 1a). Closer inspection by Stimulated Emission Depletion (STED) microscopy showed that the CD31 labelling had a speckled appearance that again was more concentrated at cell junctions (Figure 1b). Piezo1 labelling displayed as larger spots that were indicative of trimeric Piezo1 channels^23^ and again concentrated at cell junctions (Figure 1c). Merger of the CD31 and Piezo1 images exposed distinct and common localisations consistent with proximity of less than 10 nm in some instances (Figure 1d). The data suggest close co-localisation of CD31 and Piezo1 particularly at cell-cell junctions.

**Figure 1.**
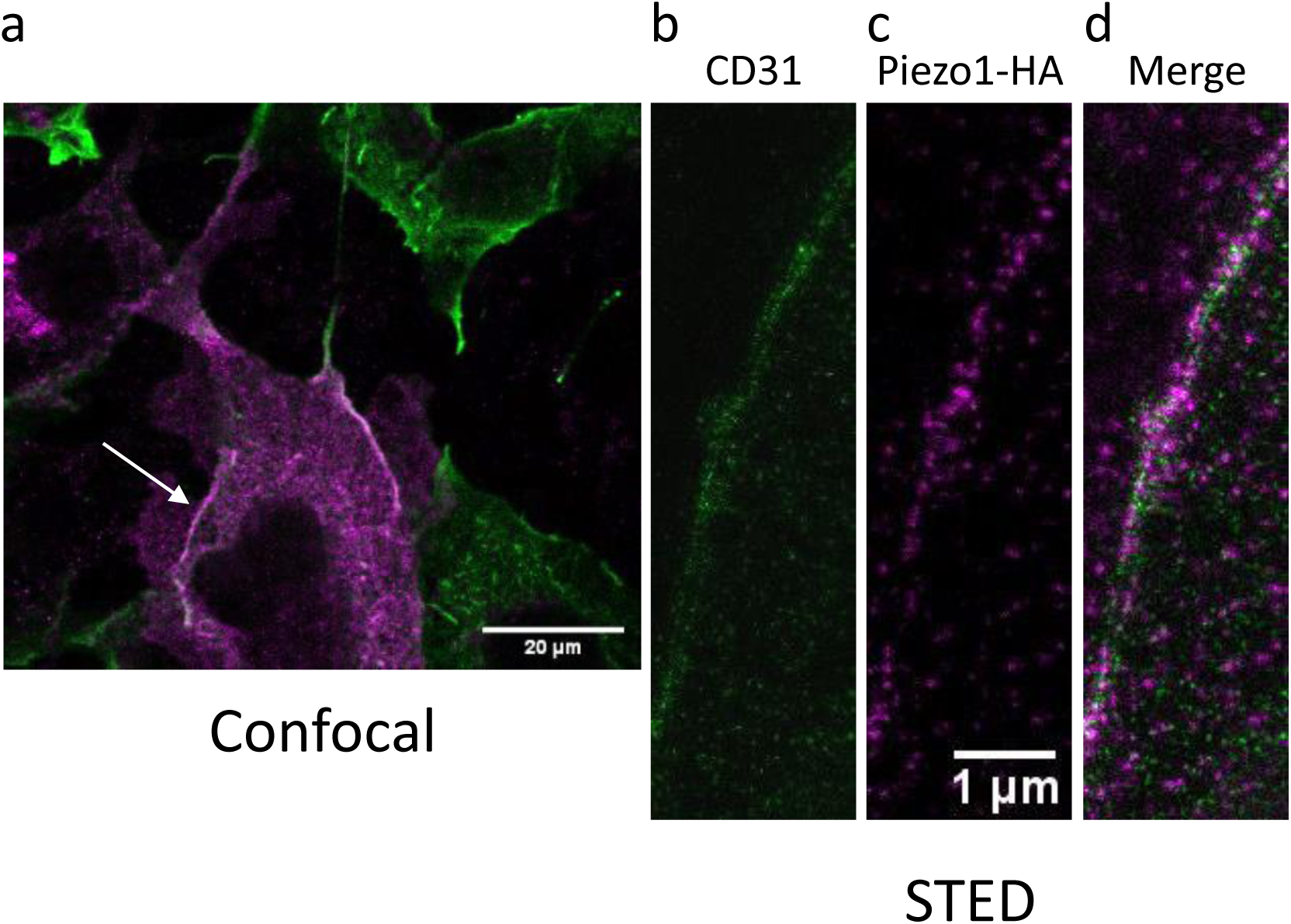
Piezo1 co-localises with CD31. COS-7 cells expressing the indicated constructs were studied by confocal microscopy (**a**) or STED microscopy (**b**-**d**). **a**) Merged images for CD31 (green) and HA-tagged Piezo1 (magenta) immunostained with anti CD31 (extracellular domain) and anti HA (extracellular tag) in unpermeabilised cells. The CD31 was tagged with SYFP2, the fluorescence of which is not displayed in the figure. The arrow indicates the region selected for STED. **b**) STED image of CD31, **c**) STED image of Piezo1-HA, **d**) merged CD31 (green) and Piezo1 (magenta) STED images with areas of overlap between the two channels indicated in white. Representative images from 3 independent experiments.

We therefore investigated if protein interactions occur, using fluorescence lifetime imaging microscopy (FLIM) to quantify Förster Resonance Energy Transfer (FRET). To this end we engineered a donor fluorophore (mTurquoise2) on the C-terminus of Piezo1 and an acceptor fluorophore (SYFP2 or mVenus) on the C-terminus of CD31 (SYFP2), VE-cadherin (mVenus) and VEGFR2 (SYFP2), enabling investigation of cytoplasmic interactions. Expressed alone, Piezo1-mTurquoise2 was primarily perinuclear and endoplasmic reticular in localisation (Figure 2a). Its lifetime map showed a single distribution with peak at about 4 ns (Figure 2a). Co-expression with CD31-SYFP2 caused Piezo1-mTurquoise2 enrichment at areas of cell-to-cell contact and lower lifetime values, consistent with FRET and thus interaction (Figure 2b, e). There was a similar result when Piezo1-mTurquoise2 was co-expressed with VE-cadherin-mVenus (Figure 2c, f) but not when it was co-expressed with VEGFR2-SYFP2, which failed to interact or concentrate Piezo1 at cell junctions (Figure 2d, g). Piezo1 interaction with CD31 was similar at intracellular sites and cell junctions whereas VE-cadherin interaction was apparently greater at junctions, although there was insufficient power for statistical significance (Supplementary Figure 1). The data suggest that CD31 and VE-cadherin promote localisation of Piezo1 to junctions and that they physically interact with Piezo1.

**Figure 2.**
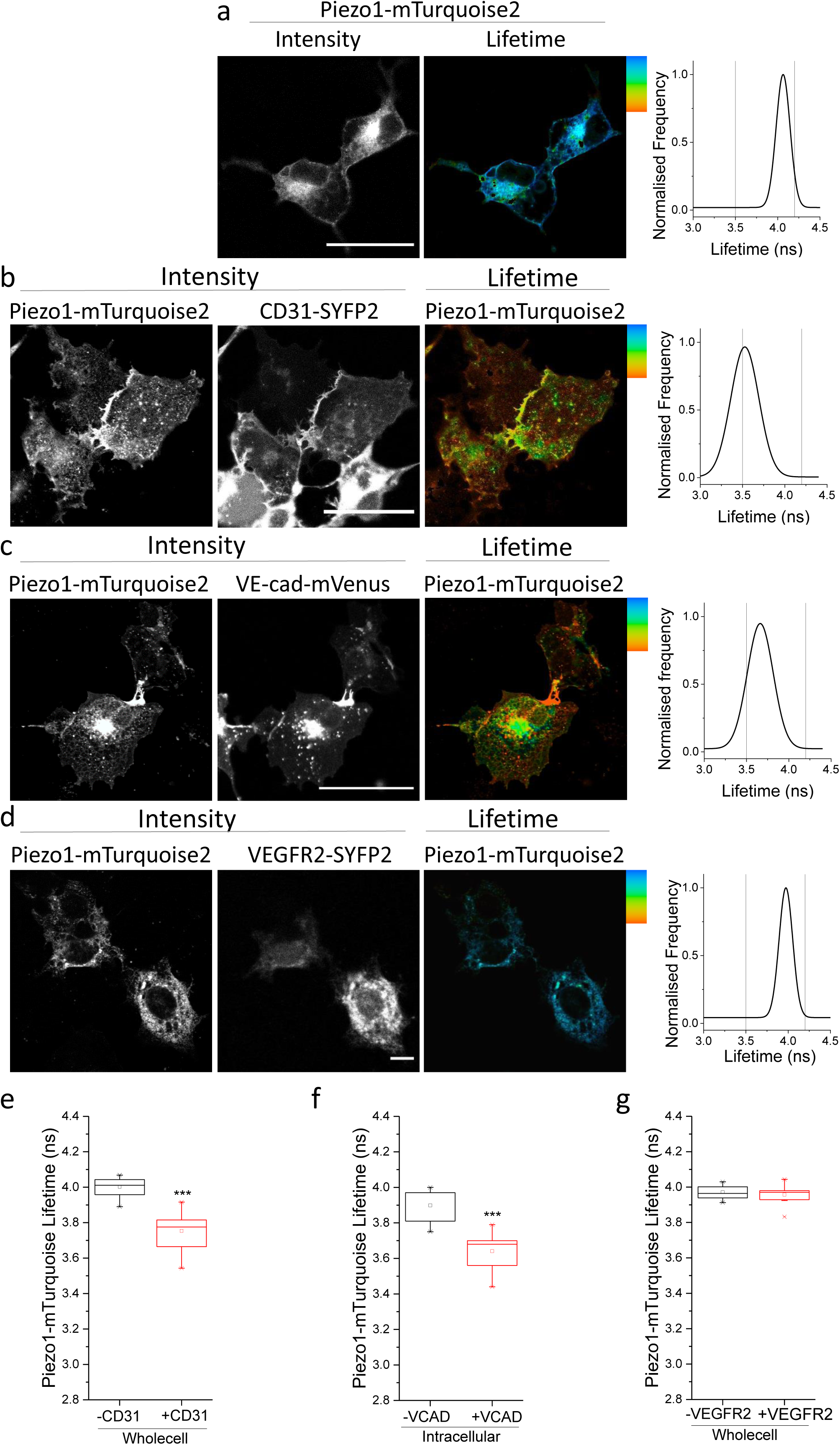
Piezo1 interacts with CD31 and VE-cadherin. COS-7 cells expressing the indicated constructs were studied by FRET/FLIM. **a**) Intensity images (left, black and white) and lifetime map (right, rainbow coloured) of Piezo1-mTurquoise2 alone. The plot on the far right is the lifetime distribution. The impact of co-expression of CD31-SYFP2 (**b**), VE-cadherin-mVenus (**c**) or VEGFR2-SYFP2 (**d**) on the intensity and lifetime distribution of Piezo1-mTurquoise2. In **a**-**d** the scale bars are 50 μm and the range represented by the rainbow colour bar is 3.5 to 4.2 ns. **e**-**g**) Mean ± s.e.mean lifetimes calculated for n=3 (VE-cadherin and VEGFR2) and n=4 (CD31) independent repeats. Changes in mean ± s.e.mean lifetime over the whole image are plotted for (**e**) Piezo1/CD31 (P = 3.16×10^−5^), (**f**) Piezo1/VE-cadherin (P = 7.68×10^−4^) and (**g**) Piezo1/VEGFR2 (P>0.05).

To determine if mechanical force affects the sub cellular localisation or interaction we applied preconditioning fluid flow and then, after a static period, a second short pulse of flow. The preconditioning tended to increase junctional relative to intracellular interaction of Piezo1 with CD31 but the second pulse of flow had no further effect (Figure 3a-d). More striking was that preconditioning enhanced junctional relative to intracellular interaction of Piezo1 with VE-cadherin and that the second pulse of flow further increased the junctional interaction (Figure 3e-g). There was a hint that flow caused mild junctional interaction between Piezo1 and VEGFR2 but statistical significance was not achieved (Figure 3h-j). The data suggest that elements of the interaction can be force sensitive.

**Figure 3.**
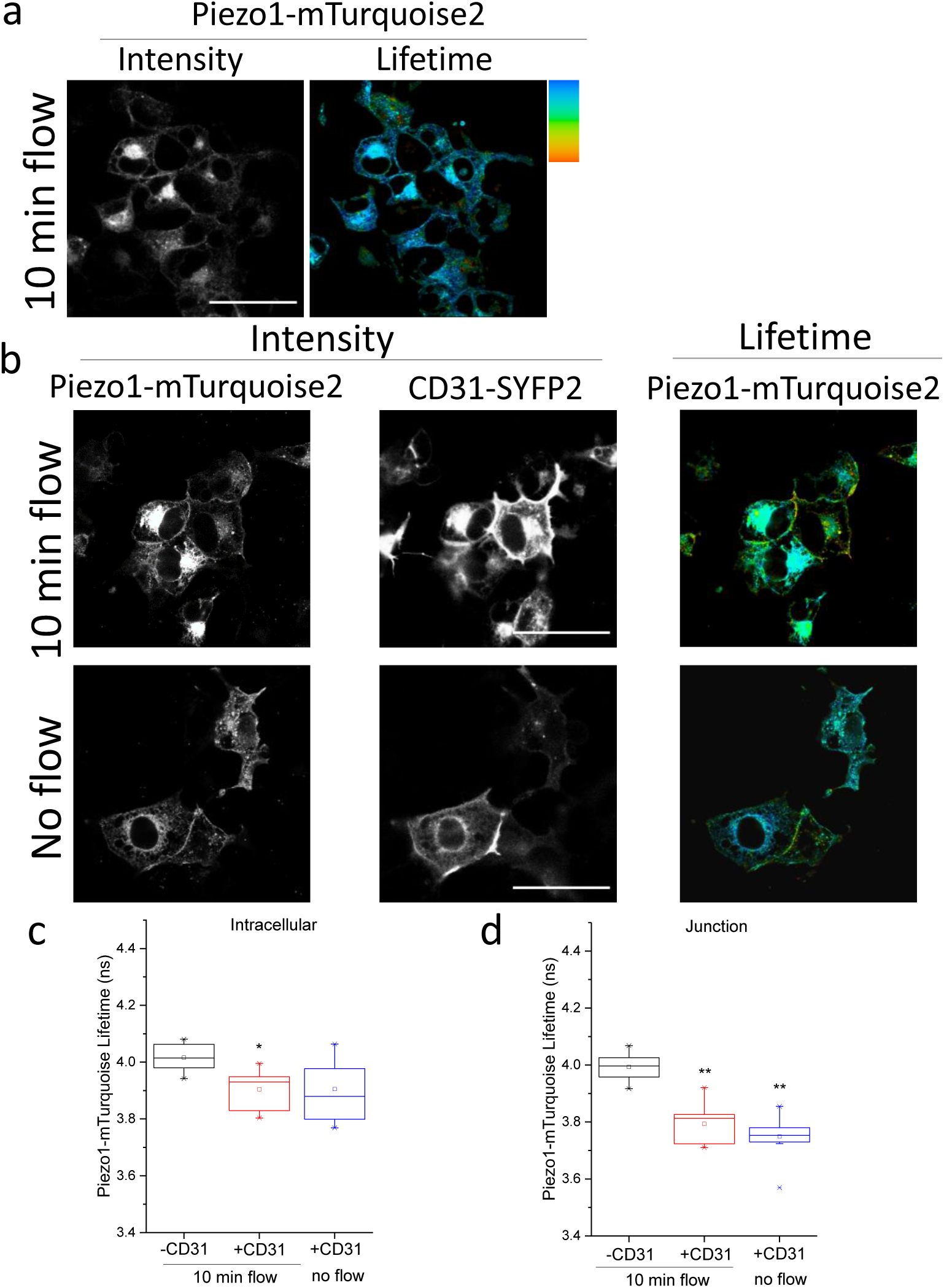
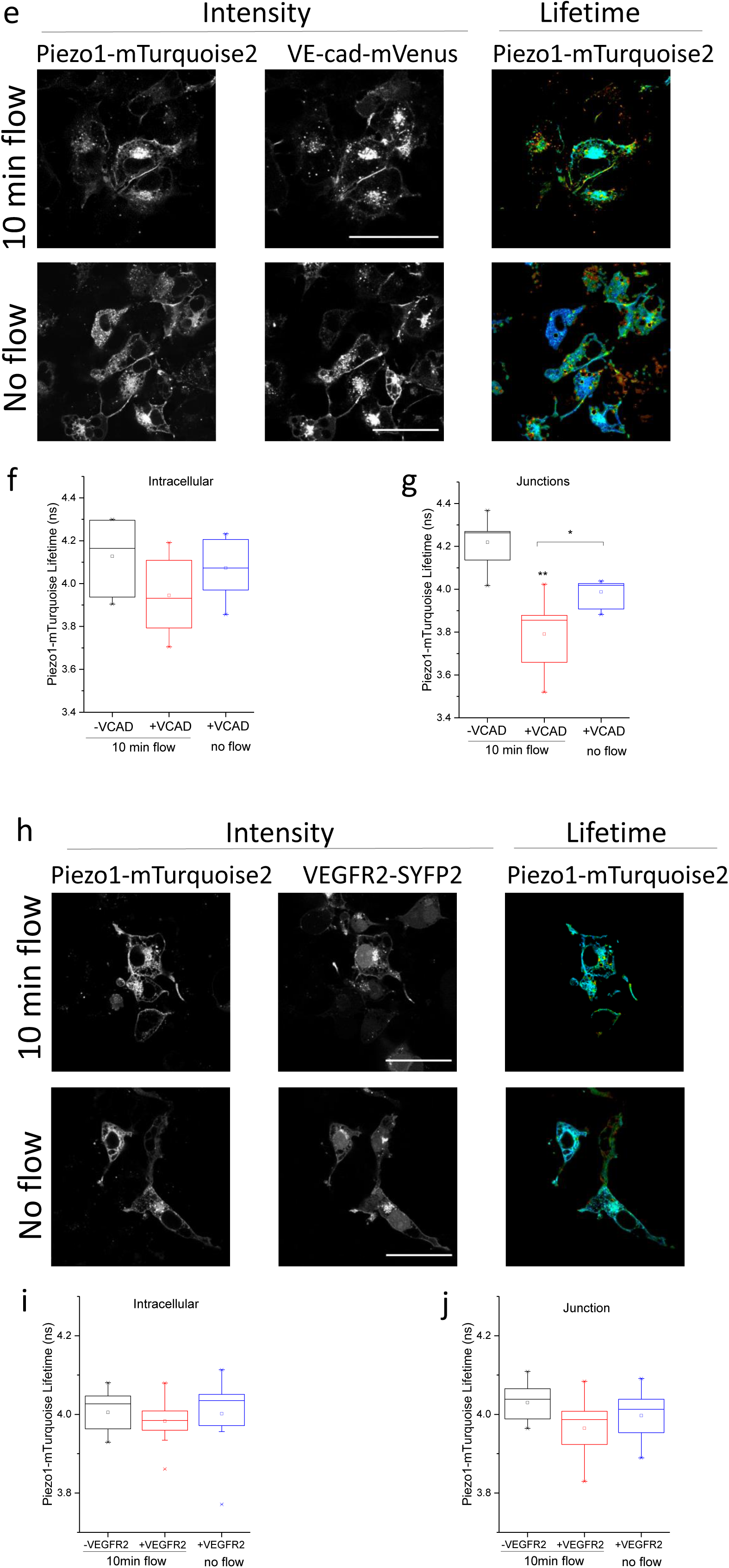
Interaction is sensitive to mechanical force. COS-7 cells were studied by FRET/FLIM after expressing Piezo1-mTurquoise2 (**a**) with or without CD31-SYFP2 (**b**-**d**), VE-cadherin-mVenus (**e**-**g**) or VEGFR2-SYFP2 (**h**-**j**). Prior to imaging, cells were preconditioned by 24-hr exposure to laminar fluid flow (shear stress = 10 dyn.cm^−2^) followed by stopped flow for 30 min (static) and then a 10-min pulse of repeated flow or no flow. **a**) Intensity (left) and lifetime (right) images of Piezo1-mTurquoise2 under 10 min flow conditions. **b**) Intensity images of Piezo1-mTurquoise2 (left) and CD31-SYFP2 (middle) and lifetime image of Piezo1-mTurquoise2 (right). Plots show the mean ± s.e.mean lifetimes of Piezo1-mTurquoise2/CD31-SYFP2 under 10 min flow or no flow at **c**) intracellular regions (P=0.02, 10 min flow) or **d**) cell-cell junctions (P=0.02, 10 min flow; P=0.01 no flow). **e**) Intensity images of Piezo1-mTurquoise2 (left) and VE-cadherin-mVenus (middle) and lifetime image of Piezo1-mTurquoise2 (right). The mean ± s.e.mean lifetimes of Piezo1-mTurquoise2/VE-cadherin-mVenus under 10 min flow or no flow at **f**) intracellular regions (P>0.05) or **g**) cell-cell junctions (P=0.002, 10 min flow; P=0.03, no flow). **h**) Intensity images of Piezo1-mTurquoise2 (left) and VEGFR2-SYFP2 (middle) and lifetime image of Piezo1-mTurquoise2 (right). The mean ± s.e.mean lifetimes of Piezo1-mTurquoise2/VEGFR2-SYFP2 under 10 min flow or no flow at **i**) intracellular regions (P>0.05) or **j**) cell-cell junctions. Mean ± s.e.mean lifetimes were calculated for n=3 independent repeats in all cases. In **a**, **b**, **e** and **h** the scale bars are 50 μm and the range represented by the rainbow colour bar is 3.5 to 4.2 ns

To further test our hypothesis that cell adhesion molecules interact with Piezo1 we focussed on CD31. Based on previously suggested and predicted functional amino acids in CD31 we selected five C-terminal residues to mutate and investigated the consequences (Figure 4a). The interaction was unchanged or potentially slightly reduced by C622A, Y663F, Y690F or S700F (Figure 4b,c *cf* Figure 2e). Most convincing, however, was that the distal C-terminal mutation Y713F prevented interaction, notably at cell junctions (Figure 4b, c). Importantly, CD31-Y713F was expressed and localised normally compared with wildtype CD31 (Figure 4d). The data support the conclusion that CD31 interacts with Piezo1 by showing that mutation of the single residue Y713F is sufficient to prevent FRET. The data also suggest that interaction occurs intracellularly and that it might be regulated by tyrosine phosphorylation.

**Figure 4.**
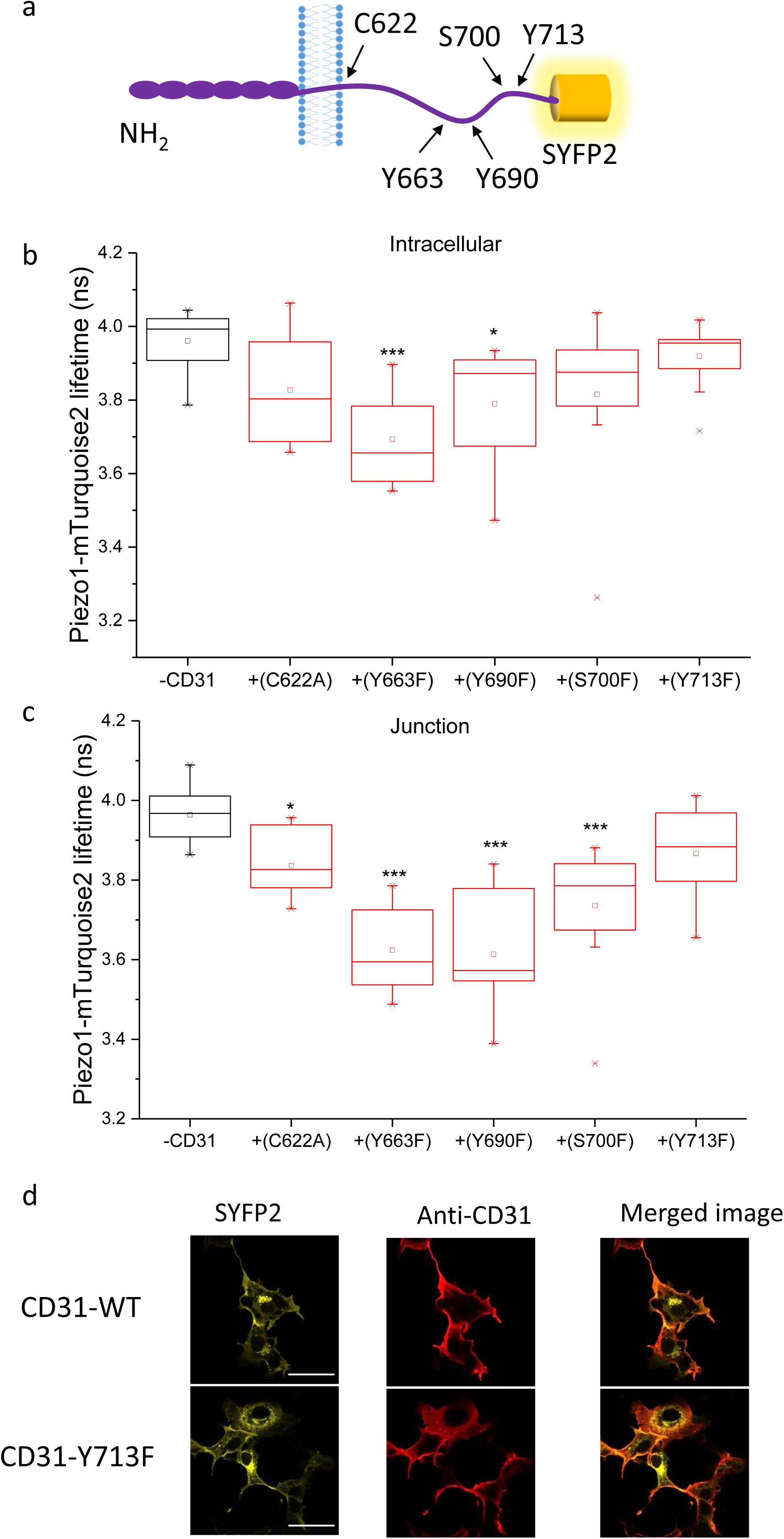
Intracellular tyrosine residue Y713 is critical for interaction. Data are shown for COS-7 cells expressing Piezo1-mTurquoise2 with or without wildtype or mutant CD31-SYFP2. **a**) Schematic of CD31 illustrating the 5 mutated residues. The interaction of Piezo1-mTurquoise2 with CD31 mutants was measured by FRET/FLIM in the **b**) intracellular regions (P=5.98×10^−4^, Y663F; P=0.02 Y690F) and **c**) cell-cell junctions (P=0.014, P=3.56×10^−4^, P=7.76×10^−4^, P=2.07×10^−4^). Mean±s.e.mean lifetime values were calculated over n=3 independent repeats. **d**) Surface expression determined by confocal imaging after immunostaining of the CD31 extracellular domain in unpermeabilised cells. Fluorescence from SYFP2 is also shown to indicate the total CD31. Scale bars are 50 μm and the range represented by the rainbow colour bar is 3.5 to 4.2 ns. Representative images are from n=3 independent experiments.

We next considered the N-terminal extracellular domain by replacing the transmembrane and cytoplasmic regions of CD31 with a plasma membrane-targeting sequence and locating SYFP2 to the intracellular site to generate CD31-ex-SYFP2 (Figure 5a). Intensity images of Piezo1-mTurquoise2 and CD31-ex-SYFP2 showed pronounced enrichment at cell junctions (Figure 5b). Lifetime analysis showed strong interaction at junctions but no interaction at intracellular sites (Figure 5c-e). The data suggest that the extracellular N-terminus of CD31 also interacts with Piezo1 channels and that this interaction is blocked prior to the CD31 reaching junctional sites.

**Figure 5.**
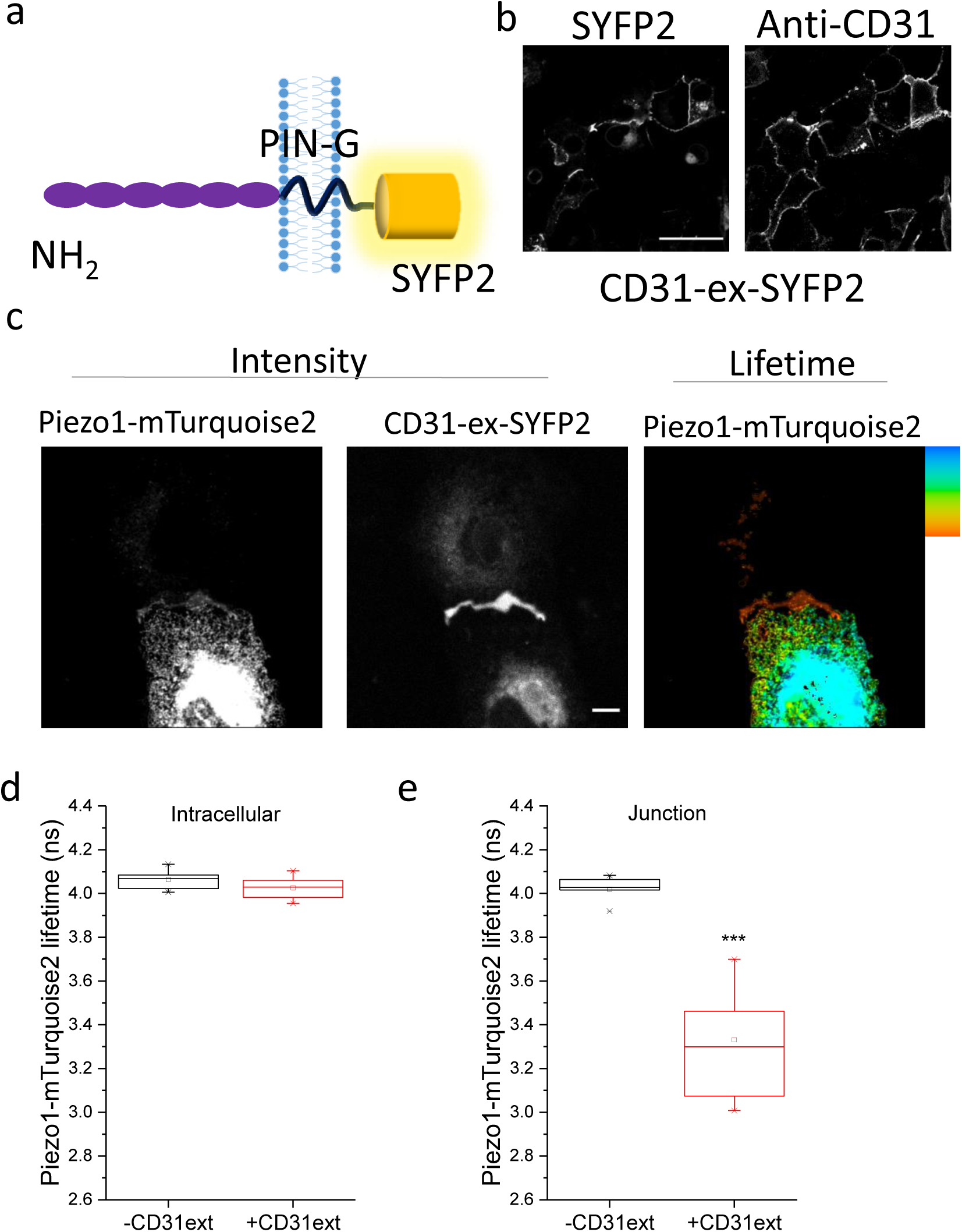
Extracellular CD31 interacts at cell junctions. Data are shown for COS-7 cells expressing Piezo1-mTurquoise2 with and without PIN-G-tagged N-terminal CD31-ex-SYFP2. a) Schematic of CD31-ex-SYFP2. **b**) Surface expression of CD31-ex-SYFP2 determined by confocal imaging after immunostaining of the CD31 extracellular domain with anti-CD31 antibody. Cells were unpermeabilised. Fluorescence from SYFP2 is also shown, reflecting total CD31. Scale bar 50 μm. Representative images from n=3 independent experiments. **c**) Intensity images and lifetime image of Piezo1-mTurquoise2 showing enrichment of CD31-ex-SYFP2 at points of cell-to-cell contact, where there were also lower lifetime values. The scale bars are 50 μm and the range represented by the colour bar is 3.5 to 4.2 ns. Mean ± s.e.mean lifetime values for Piezo1-mTurquoise only and in the presence of CD31-ex-SYFP2 at the **d**) intracellular regions and **e**) cell-cell junctions (P=4.12×10^−4^).

To investigate if the interaction has functional significance we measured mechanical sensitivity of Piezo1 channels in excised outside-out patches, thus restricting data collection to the surface membrane. Positive pressure steps caused membrane stretch, which activated Piezo1 channels (e.g. Figure 6a). CD31 shifted the pressure-response curve to the right and reduced its slope (Figure 6b). To determine how CD31 might reduce the mechanical sensitivity of Piezo1 channels we made C-terminally truncated Piezo1 constructs (Figure 6c) and tested their ability to interact with CD31 by co-immunoprecipitation. We found that N-terminal regions of Piezo1, lacking the C-terminal ion pore and CED, were sufficient for interaction (Figure 6d). The data suggest that CD31 interacts with the N-terminal propeller blade structures of Piezo1 channels^23–25^ to potentially alter their flexibility and tone down channel response to increases in force.

**Figure 6.**
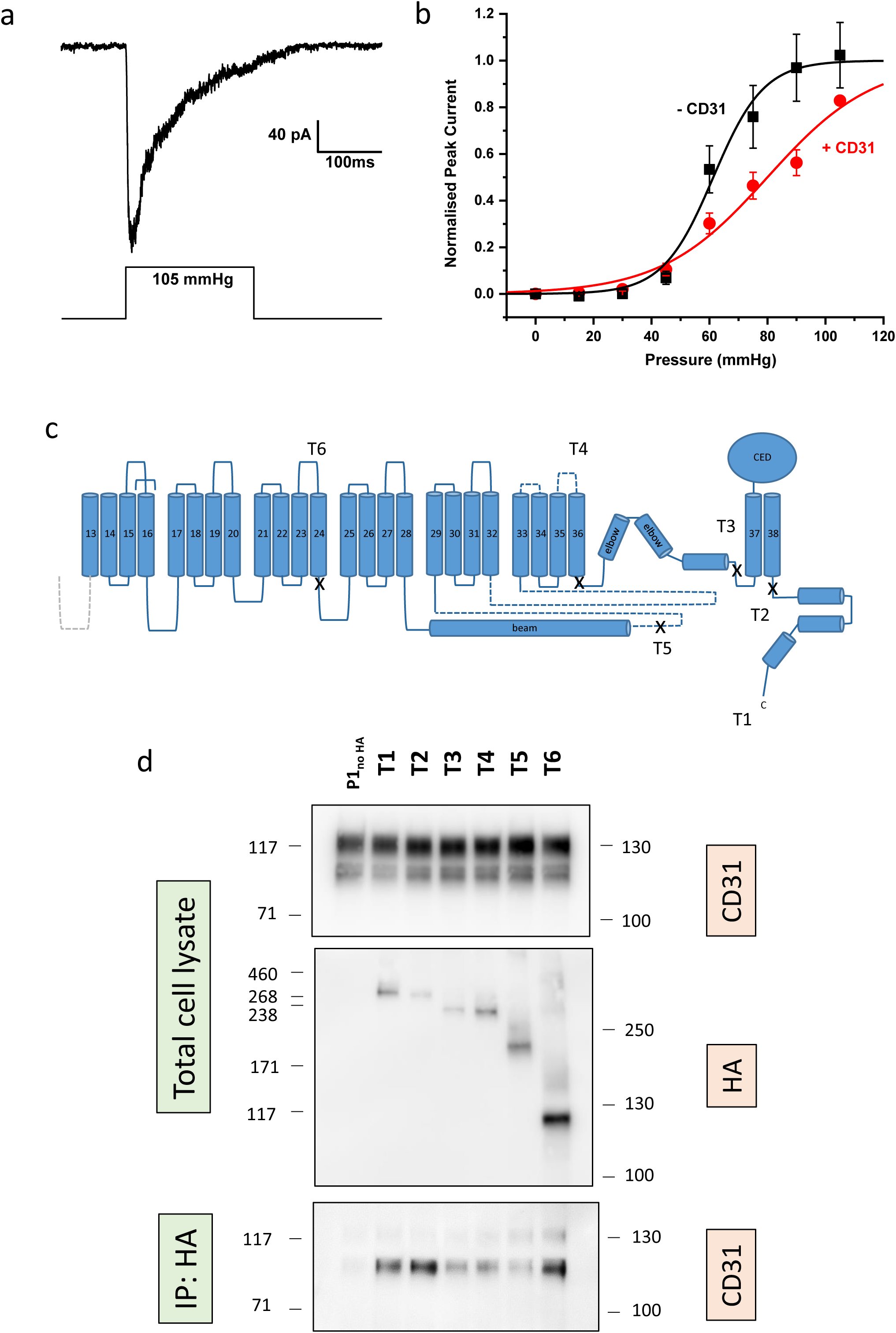
CD31 reduces mechanical sensitivity of Piezo1 channels. Data are shown for T-REx-293 cells containing stable inducible expression of Piezo1 with or without transient transfection with SYFP2 or CD31-SYFP2 (**a**, **b**) or HEK 293 cells stably-expressing wildtype CD31 and transiently expressing HA-tagged Piezo1 or HA-tagged Piezo1 fragments (**d**). Representative current trace for an outside-out patch recording from a Piezo1 plus SYFP2 cell (minus CD31, - CD31). Currents were evoked by a series of positive pressure pulses at a constant holding voltage of −80 mV; the example is for a pressure step to 105 mmHg from 0 mmHg. **b**) Analysis of experiments of the type exemplified in (**a**) showing peak current plotted against the amplitude of the pressure pulse. The fitted curves are the fitted Boltzmann equation giving mid-point (P_50_) and slope values of 61.3 mmHg and 8.70 for Piezo1 plus SYFP2 (n=9) and 79.99 mmHg and 18.31 for Piezo1 plus CD31-SYFP2 (n=9). **c**) Schematic of full-length Piezo1 (T1) and the positions of the truncations to create the Piezo1 fragments T2-T6. The proximal N-terminus (membrane-spanning segments 1-12) is not illustrated. **d**) Typical blots for wildtype Piezo1 (P1_noHA_) and HA-tagged T1-T6 Piezo1 expressed in the stable CD31 cells, showing total cell lysate and CD31 detected by anti-CD31 antibody in the anti-HA immunoprecipitate (IP: HA). CD31 was detected in all IPs except, as expected, when Piezo1 lacked the HA tag (P1_noHA_). Mature and immature CD31 were detected in total cell lysate and IP but the immature protein was more prominent in the IP.

## Discussion

The study suggests that two apparently divergent ideas for endothelial flow sensing - Piezo1 and the CD31 triad - are interrelated. We show evidence of physical proximity detected by super-resolution microscopy techniques and of physical interaction detected by FRET, co-immunoprecipitation and functional effect of over-expressed CD31 on activity of Piezo1 channels in excised membrane patches. Intriguingly we found that the second cell adhesion molecule of the triad - VE-cadherin - also interacts with Piezo1 (in the absence of CD31) and that its interaction has distinctive flow sensitivity. VEGFR2, which is recruited to the triad in response to flow^11^, did not directly interact with Piezo1 as determined by FRET; we hypothesize that it may depend on CD31 or VE-cadherin for efficient association.

We show that one consequence of Piezo1-CD31 interaction is the reduced mechanical sensitivity of Piezo1 channels characterized by rightward shift in the pressure-response curve and shallower slope, suggesting that, the more Piezo1 channels interact with CD31, the more their force sensitivity is tempered. This effect may be needed because increased force associated with cell-to-cell contact would otherwise over-activate Piezo1 channels and cause too much local cytoplasmic Ca^2+^ elevation. Appropriate Ca^2+^ elevation is important for numerous proteins, including CD31 because of its suggested calpain-dependent cleavage^30^ and VE-cadherin because of its inherent Ca^2+^ sensitivity^14,15^. The importance of calpain, a Ca^2+^-activated protease, as a downstream mediator of Piezo1 effects, regulator of cytoskeletal anchorage complexes and component of the endothelial flow sensing machinery has been proposed^17,31–33^. Therefore the functional relationship between Piezo1 and CD31 is likely to be two-way, including but not necessarily restricted to: physical interaction of CD31 with Piezo1 channels to alter channel sensitivity; and Ca^2+^ entry through Piezo1 channels to activate calpain and therefore modify CD31 and other proteins.

The CD31 triad and its relationship to Piezo1 channels occurs at cell-cell junctions that are likely to be physically hidden from direct impact of fluid flow. This is not to say that they lack importance as mediators of the flow response but the luminal surface of endothelial cells is the more obvious site for impact of fluid flow either directly or via glycocalyx. While the functional significance of CD31 may be restricted to points of cell-to-cell contact, we know that flow sensing Piezo1 channels also exist at the luminal surface because patches of membrane picked from this surface of native endothelium contain such channels^18^. These channels may not be associated with CD31 and could therefore be expected to achieve greater mechanical sensitivity.

Overall the data suggest previously unrecognised physical interaction, co-localisation and functional relationship between the Piezo1 force sensing ion channel and cell adhesion molecules. The evidence brings together what had appeared to be contradictory observations on flow sensing, suggesting partnership at cell junctions but potentially independent functions, at least of Piezo1, at the luminal membrane. The relationship of Piezo1 to adhesion molecules may extend beyond CD31 and VE-cadherin to other cell adhesion molecules, which are numerous in endothelial cells and other cells types^15^. The findings potentially open up a new opportunity to better understand how force sensing is achieved and transduced at cell-cell junctions where the integration with Ca^2+^-entry channels could have high importance. While Piezo1 and CD31/VE-cadherin are important in responses to fluid flow, they also have other roles, most notably in regulating junctional integrity and thus permeability and tissue structure that depend on cells detecting and responding appropriately to forces between cells.

## Supplementary Information

is provided as a separate document.

## Acknowledgements

The study was supported by funding from the Wellcome Trust and British Heart Foundation. We thank Sally Boxall (Leeds Bio-Imaging Facility) and David Myers for microscopy technical support and Bing Hou, Sarka Tumova, Claudia Bauer and Deborah Linley for data that encouraged elements of the study but which are not included in this report.

## Author Contributions

EC-B and MJL designed protein constructs. EC-B, MJL and HJG cloned protein constructs. EC-B designed and performed microscopy experiments. OVP set up and performed pressure patch clamp experiments. MJL designed and performed immunoprecipitation experiments. EC-B, OVP, MJL and DJB analysed data. PDB guided statistical analysis of FRET/FLIM data. EC-B and DJB directed the study, made the figures and wrote the manuscript. DJB conceptualised the study and supervised the project team.

## Methods

### Cell lines

COS-7 cells were from the American Type Culture Collection (ATCC) and maintained in Dulbecco’s Modified Eagle Media (DMEM) supplemented with 10% FCS, 2 mM L glutamine, 100 U.ml^−1^ penicillin, 100 µg.ml^−1^ streptomycin in 5% CO_2_, 95% air atmosphere. Cells were sub-cultured upon reaching a surface area density of 80–90% by detaching with 0.5% trypsin. T-REx-293 cells from Thermo Fisher Scientific were conferred with tetracycline inducible overexpression of human Piezo1^18^ were maintained in DMEM supplemented with 10% FCS, 2 mM L glutamine,100 U.ml^−1^ penicillin, 100 µg.ml^−1^ streptomycin in 5% CO_2_, 95% air atmosphere. Cells were selected with zeocin (400 μg.ml^−1^) and blasticidin (5 μg.ml^−1^) and induced with tetracycline (100 ng.ml^−1^) for 24 hr. T-REx-293 overexpressing CD31 were generated by seeding at 70% confluency in a 6 well plate. Cells were transfected for 5 hours with 500 ng CD31 plasmid and 0.3% Lipofectamine 2000 (Invitrogen). After 48 hr selection with 5 μg/ml blasticidin began. Single cell lines stably expressing CD31 were isolated.

### Constructs and cloning

human Piezo1-mTurquoise2 was sub-cloned from human Piezo1-GFP^17^. pSYFP2-C1^34^ (Addgene plasmid # 22878; http://n2t.net/addgene:22878; RRID:Addgene_22878) was a gift from Dorus Gadella. mTurquoise2-C1^35^ (Addgene plasmid # 54842; http://n2t.net/addgene:54842; RRID:Addgene_54842) was a gift from Michael Davidson and Dorus Gadella. Human CD31 was obtained from Origene (TrueClone, SC119894). Piezo1 and CD31 pcDNA6 templates were generated by inverse PCR with the Phusion® DNA polymerase (New England Bio Labs). mTurquoise2 and SYFP2 were incorporated to the Piezo1/CD31 templates by overlap PCR using In-Fusion® HD cloning kit (Clonetech). Linkers were then attached between Piezo1/mTurquoise2 and CD31/SYFP2 using In-Fusion® HD cloning kit (Clonetech). For Piezo1/mTurquoise2 the linker was flanked by a BamH1 restriction. The linker on CD31/SYFP2 was flanked by a HindIII restriction site. VEGFR2-SYFP2 was sub-cloned from mEmerald-VEGFR2-N1 (Addgene plasmid # 54298; http://n2t.net/addgene:54298; RRID:Addgene_54298) was a gift from Michael Davidson. When sequenced mEmerald-VEGFR2-N1 was found to contain a frameshifting CT insertion between residues T1303-A1304 and the single nucleotide polymorphisms (SNP) V297I and H472Q. Site-directed mutagenesis was performed to remove the insertion and correct the SNPs. To facilitate cloning of VEGFR2-SYFP2, a linker flanked by AgeI and SacII restriction sites was introduced into pcDNA™4/TO between EcoRI and XhoI restriction sites using Gibson Assembly® (New England Biolabs, Ipswich, MA, USA). The SYFP2 fluorophore was inserted downstream of the linker between SacII and XbaI restriction sites using pSYFP2-C1. PCR products were assembled using Gibson Assembly® to insert VEGFR2 upstream of the linker. The C-terminal fluorophore was removed from this construct using Gibson Assembly® to assemble VEGFR2 and vector PCR products. mVenus-VE-Cadherin-N-10 (Addgene plasmid # 56340; http://n2t.net/addgene:56340; RRID:Addgene_56340) was a gift from Michael Davidson and was used as acquired. To generate C-terminal HA-tagged Piezo1 the C-terminal GFP from the human Piezo1-GFP construct^17^ was first removed by ligating the KpnI and NotI restriction enzyme digested vector and Piezo1 PCR product. This construct was subsequently used as a template to create full-length and truncated Piezo1 expression vectors. PCR products covering (T1) full length Piezo1 sequence to L2471, G2174, I2089, S1591 and A1128 were produced and inserted into the vector using Gibson Assembly®. Piezo1-HA (CED). An HA-tag was also introduced between T2413 and C2414 located in the C-terminal extracellular domain. CD31 mutants were generated using the CD31-SYFP2 plasmid as a template. Mutagenesis was carried out using PrimeSTAR HS DNA polymerase (TaKaRa). The extracellular domain of CD31 containing SYFP2 and PIN-G was cloned from the CD31-SYFP2 plasmid using In-Fusion® HD cloning kit (Clonetech).

### Paraformaldehyde fixation

Cells were fixed with 4% paraformaldehyde (10 min, RT) and washed with PBS 3 times for 5 min. Fixation was quenched with glycine (0.1 M in PBS, 10 min, RT) and cells were washed with PBS (3 times, 5 min, RT). Fixed cells were covered in PBS and stored at 4°C. All fixation steps were carried out under low light and sterile condition. Samples were imaged on the same day or stored for a maximum of 24 hr.

### STED imaging

COS-7 cells were grown on coverslips placed inside 12-well plates. Wells were seeded with ~ 1 × 10^6^ cells in 3 ml cell culture medium. Cells were transfected using FuGene with Piezo1-HA and CD31-SYFP2 per well. After 48hr cells were fixed and transferred to 12 well plates blocked with 1% BSA (in PBS) (15 min at RT). Primary antibodies against HA (rabbit anti-HA, Cell Signaling Technology, #3724, 1:800) and CD31 (mouse anti-CD31, 1:200) were diluted in 1% BSA/PBS and added to cells for 1 hr at 37°C. After incubation cells were washed 3 times (5 min, RT) with PBS. Secondary antibodies against rabbit (anti-rabbit 580, Aberrior, 41367, 1:100) and mouse (anti-mouse STAR red, Aberrior,52283,1:100) diluted in PBS were added to the cells and incubated (30 min, RT). Cells were washed 3 times with PBS (5 min, RT) and mounted using ProLong® Gold antifade mountant (ThermoFisher, P36930). STED microscopy was performed using a 100× objective on the STEDYCON 2-colour STED imaging system (Abberior Instruments). Images were exported to Fiji^36^ for final processing and assembly.

### FLIM sample preparation

Cells imaged under static conditions 35-mm plastic cell culture dishes were plated with ~ 3 × 10^6^ cells in 3 ml cell culture medium. After subculture cells were transfected using FuGene with Piezo1-mTurquoise. For co-transfections acceptor labelled constructs were added. After 48 hr cells were fixed with 4% paraformaldehyde. For flow experiments cells (~ 1 × 10^6^ in 200 µl medium) were plated onto the ibiTreat µSlide-I^0.8^ Luer (Ibidi) and allowed to attach for 24 hr. Transfections were carried out using Lipofectamine™ 3000. The medium in each slide was replaced with the transfection mixture in fresh medium and cells were incubated for 24 hr. To culture and stimulate cells under flow conditions we used a system comprising an air pressure pump, pump control software, perfusion set (yellow/green) and fluidics unit (Ibidi). Cells were exposed to shear stress of 10 dyn.cm^−2^ for 24 hr. Flow was stopped for a 30 min rest period after which cells were stimulated with flow (10 dyn.cm^−2^, 10 min) and then fixed with 4% paraformaldehyde.

### FLIM microscopy

Intensity and FLIM images were obtained on an upright LSM710 (Carl Zeiss) microscope with a 40× /1.0 NA, water-dipping objective or 63×/1.40 Oil (Carl Zeiss). Acceptor intensity images were obtained with excitation at 512 nm using an Argon laser and registered on the Zeiss PMT detectors. Two-photon excitation was provided by Chameleon (Coherent) Ti:Sapphire laser tuned to 800 nm. FLIM emission events were recorded by an external detector (HPM-100, Becker & Hickl) attached to a commercial time-correlated single photon counting electronics module (Becker & Hickl) with a 480/40 (Chroma) emission filter. FLIM images were fitted using in SPCImage (Becker &Hickl). A single component incomplete multi-exponential model was used with a laser repetition time of 12.5 ns. Colour-coded lifetime maps and grayscale intensity images were exported from SPCImage. Acceptor intensity images were processed using Fiji. The histogram intensity weighted mean lifetimes for each image was generated by SPCImage 5.6; values were exported to OriginPro. The peak values were obtained by doing a Gaussian fit.

### Immunostaining

COS-7 cells (~ 1 × 10^6^ /ml medium, 30 µl) were plated onto ibiTreat µSlide-VI^0.4^ Luer (Ibidi) and allowed to attach for 24 hr. Transfections were carried out using Lipofectamine™ 3000. After 48 hr cells were fixed and blocked with 1% BSA (in PBS) for 15 min at RT. Primary antibodies against CD31 (mouse anti-CD31, clone JC70A, Dako1:200) were diluted in 1% BSA/PBS and added to cells for 1 hr at 37°C. After incubation cells were washed 3 times (5 min) with PBS. Secondary antibodies mouse (anti-mouse Alexa 594, Jackson Immuno Research, 1:300) diluted in PBS were added to the cells and incubated for 30 min at RT. Cells were washed 3 times with PBS (5 min) and stored in PBS at 4°C. Samples were imaged on the same day or stored for a maximum of 24 hr. Imaging was carried out on LSM710 (Carl Zeiss Ltd.) using a 40×/1.3 oil objective. Images were exported to Fiji for final processing and assembly.

### Patch clamp electrophysiology

T-REx-293 cells with tetracycline inducible overexpression of human Piezo1 were seeded into a T25 tissue culture flask. After 24 hr, transfections were carried out using Lipofectamine™ 3000. Cells were transfected with CD31-SYFP2 or SYFP2. Expression of Piezo1 was induced by the addition of tetracycline (100 ng.ml^−1^, 24 hr). Cells expressing Piezo1 with CD31-SYFP2 or SYFP2 were detached using trypsin and seeded onto coverslips for patch clamp experiments. Macroscopic transmembrane ionic currents through outside-out patches of T-REx-293 cells containing stable inducible expression of human Piezo1 with transient transfection with SYFP2 or CD31-SYFP2 were recorded using standard patch-clamp technique in voltage-clamp mode. Patch pipettes were fire-polished and had a resistance of 4–7MΩ when filled with pipette solution. Symmetrical Na^+^ (K^+^/Ca^2+^-free) solution of the following composition (mM): NaCl 140, HEPES 10 and EGTA 5 (pH 7.4, NaOH), was used for both, pipette and bath solutions, and the currents were recorded at −80 mV. All recordings were made with an Axopatch-200B amplifier (Axon Instruments, Inc., USA) equipped with Digidata 1550B and pClamp 10.6 software (Molecular Devices, USA) at room temperature. 200-ms pressure steps were applied directly to the patch pipette with an interval of 12 s and with an increment of 15 mmHg using High Speed Pressure Clamp HSPC-1 System (ALA Scientific Instruments, USA). Current records were filtered at 2 or 5 kHz and acquired at 5 or 20 kHz.

### Co-immunoprecipitation

500 μg of transiently transfected HEK 293 cell lysate (lysis buffer: 10 mM Tris, pH 7.4, 150 mM NaCl, 0.5 mM EDTA, 0.5% Nonidet P40 substitute, 0.1% glycerol containing protease inhibitor cocktail (Sigma)) was incubated with 1 μg anti-HA (Roche, clone 3F10) for at least 4 hr at 4°C, prior to extraction overnight using Protein G agarose (Pierce). The beads were washed three times with ice-cold lysis buffer and bound proteins eluted using sample buffer (4× SB: 250 mM Tris pH 6.8, 8% SDS, 40% glycerol, 8% β-mercaptoethanol) and heating at 95°C. Samples were loaded on 7% gels and resolved by electrophoresis. Proteins were transferred to PVDF membranes and labelled overnight with anti-HA (0.01 μg/mL, Roche clone 3F10), anti-VE-Cadherin (0.5 μg.ml^−1^, R&D Systems; MAB9381), anti-VEGFR2 (0.2 μg.ml^−1^, R&D Systems; AF357), anti-CD31 (1:1000, Dako; clone JC70A) or anti-β-actin (200 ng.ml^−1^, Santa Cruz). Horse radish peroxidase donkey anti-mouse/ anti-rat/anti-goat secondary antibodies (1:10000, Jackson ImmunoResearch) and SuperSignal Femto detection reagents (Pierce) were used for visualisation.

### Data analysis

Statistical analysis of fluorescent lifetime values was carried out using OriginPro. Normality test revealed that at least 1 dataset was not significantly drawn from a normally distributed population at the 0.05 level. Consequently, the Mann-Whitney Test was used to compare if two distributions are significantly different. For data containing 3 different conditions the Kruskal-Wallis Anova was used to check for variance, followed by use of the Mann-Whitney Test to compare data pairs. Each image was treated as an independent replicate, ordering effects were negligible. Number of experiments performed from different cell preparations are defined as n. Where multiple pairwise tests were performed on a single dataset, Bonferroni correction was applied. Patch-clamp data was acquired in a blinded experiment. The person performing the patch measurements was provided with Piezo1 expressing HEK-TRex cells transfected with either CD31-SYFP2 or SYFP2 but was not made aware of which sample contained each protein. All electrophysiological data were analysed and plotted using pClamp 10.6 and MicroCal Origin 2018 (OriginLab Corporation, USA) Software. Pressure-dependent curves were constructed in Origin and fitted with Boltzmann equation: y = A2 + (A1-A2)/(1 + exp((x-x0)/dx)).

