## Supplementary Information for "Cell adhesion molecule interaction with Piezo1 channels is a mechanism for sub cellular regulation of mechanical sensitivity"

**Supplementary Figure 1| Comparisons of intracellular and junctional localisation and interaction.** COS-7 cells expressing the indicated constructs were studied by FRET/FLIM. Intensity and lifetime images of Piezo1-mTurquoise2 with co-expression of **a**) CD31-SYFP2 and **b**) VE-cadherin-mVenus. Arrows indicate the regions of cell-cell junction and box the intracellular area. The Piezo1/CD31 lifetime distributions of the representative images are shown for the **c**, **d**) intracellular region ($P=1.26x10^{-4}$) and **e**, **f**) cell-cell junctions ($P=8.11x10^{-4}$). The Piezo1/VE-cadherin lifetime distribution of the representative image is shown for the **g**, **h**) intracellular (P=0.001) or **i**, **j**) junction regions (P=0.001) are plotted. Lifetimes were evaluated for n=3 independent repeats. The range represented by the rainbow colour bar is 3.5 to 4.2 ns.

### Supplementary Figure 1

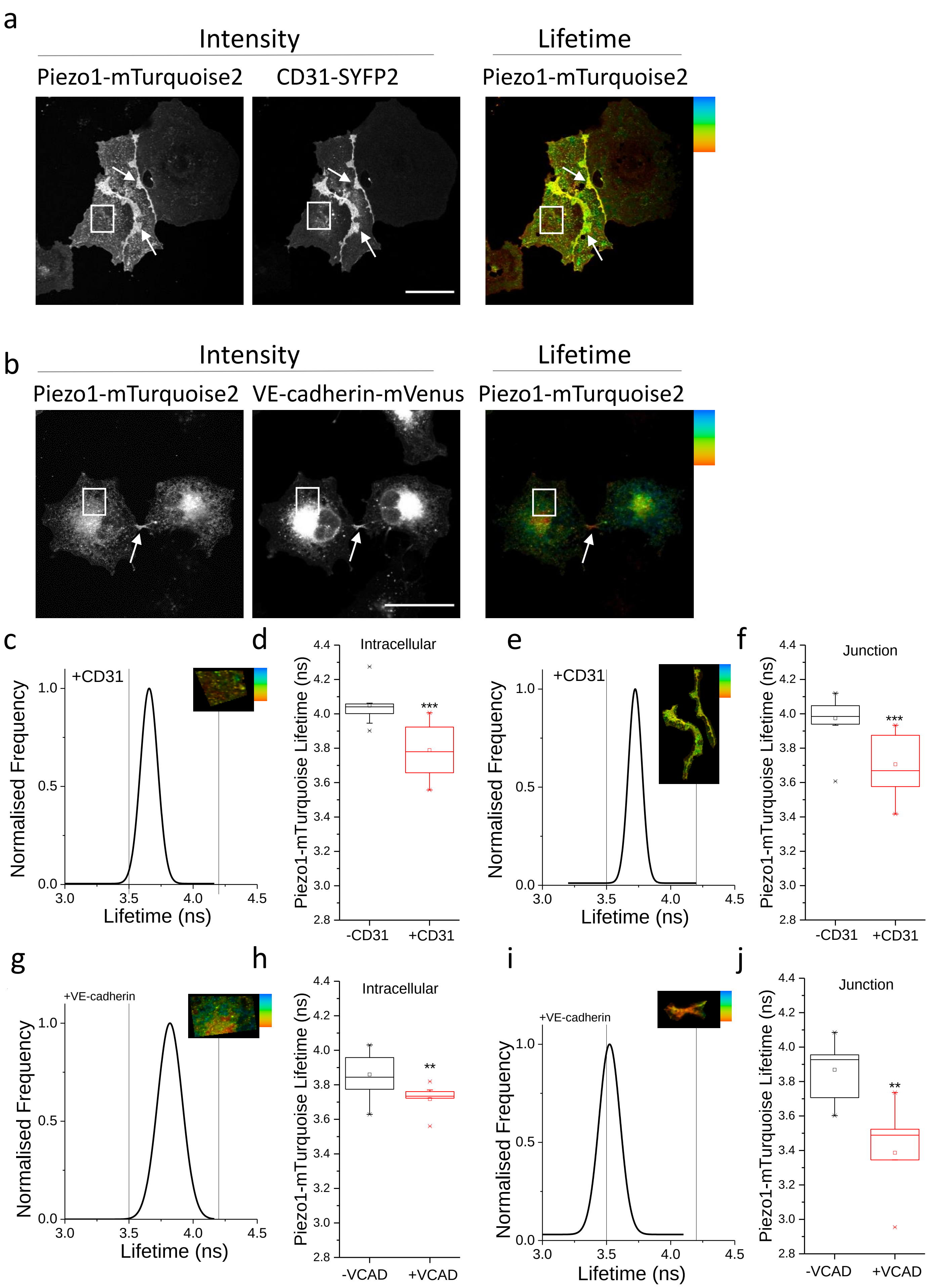
